# Identification of new marker genes from plant single-cell RNA-seq data using interpretable machine learning methods

**DOI:** 10.1101/2020.11.22.393165

**Authors:** Haidong Yan, Jiyoung Lee, Qi Song, Qi Li, John Schiefelbein, Bingyu Zhao, Song Li

## Abstract

An essential step in the analysis of single-cell RNA sequencing data is to classify specific cell types with marker genes. In this study, we have developed a machine learning pipeline called Single cell Predictive markers (SPmarker) to identify novel cell-type marker genes in the *Arabidopsis* root.

Unlike traditional approaches, our method uses interpretable machine learning methods to select marker genes. We have demonstrated that our method can (1) assign cell types based on cells that were labeled using published methods, (2) project cell types identified by trajectory analysis from one dataset to other datasets, and (3) assign cell types based on internal GFP markers.

Using SPmarker, we have identified hundreds of new marker genes that were not identified before. As compared to known marker genes, we have found more orthologous genes of these new marker genes in corresponding rice single cell clusters. We have also found 172 new marker genes for trichoblast in five non-*Arabidopsis* species, which expands number of marker genes for this cell type by 35-154%.

Our results represent a new approach to identify cell-type marker genes from scRNA-seq data and pave the way for cross-species mapping of scRNA-seq data in plants.

## Introduction

Single cell RNA sequencing (scRNA-seq) has recently emerged as a powerful approach to investigate gene expression in complex multicellular organisms. Compared to bulk RNA-seq, scRNA-seq can identify rare cell populations and reveal transitions of cell states at different developmental stages, which are difficult to capture using traditional methods (Butler et al., 2018; Trapnell, 2015; Wang and Navin, 2015). As a transformative technology, scRNA-seq is particularly important for plant research because traditional methods for determining gene expression in individual cell types rely on transgenic lines expressing cell-type-specific fluorescent markers, which are not available in most non-model species. Because of the advantages of using scRNA-seq in plants, this approach has been applied in a number of studies to profile transcriptomes of *Arabidopsis*, rice (*Oryza sativa*), tomato (*Solanum lycopersicum*), and maize (*Zea mays*) (Bezrutczyk et al., 2020; Satterlee et al., 2020). The *Arabidopsis* root is an ideal system to address important questions in plant biology using scRNA-seq, including analysis of expression pattern of rare cell types (Denyer et al., 2019; Ryu et al., 2019), determination of developmental trajectories of root cells (Denyer et al., 2019; Jean-Baptiste et al., 2019; Ryu et al., 2019; Zhang et al., 2019b), and characterization of stress responsive genes at the single cell level (Jean-Baptiste et al., 2019; Ryu et al., 2019; Shulse et al., 2019).

Identification of cell types is a key step in the analysis and interpretation of scRNA-seq data (Luecken and Theis, 2019). Currently, approaches to define *Arabidopsis* root cell types fall into three major categories: (1) ICI (index of cell identity) method. This approach uses selected marker genes based on information theoretic scores from published cell expression profiles (Efroni et al., 2015; Shulse et al., 2019; Turco et al., 2019); (2) Cluster-marker genes. This approach generates clusters of cells with unsupervised dimension reduction methods and assigns cell types by visualizing expression patterns using known marker genes (Jean-Baptiste et al., 2019; Ryu et al., 2019; Zhang et al., 2019b). (3) Correlation methods. This category of methods compute correlation coefficient between single cells (or cell clusters) and published gene expression data (Jean-Baptiste et al., 2019; Shulse et al., 2019). All these strategies rely on the knowledge of cell-type marker genes and each of these three approaches has limitations. ICI method was developed based on microarray data and only included 15 cell types with approximately 20 marker genes per cell type. However, this method has not been updated to include more data from bulk RNA-seq or single cell experiments and does not include cell types in different developmental stages. The cluster-marker gene methods used marker genes that vary from publication to publication and there is no standard to determine how many marker genes are optimal and how specific each marker gene is to a cell type. The correlation methods need expression profiles of all expressed genes from known cell types. Although the third method borrows information across many genes, one limitation is that this method might fail when assigning cell types between highly diverged species due to the loss of orthologous marker genes in non-model species (Liu et al., 2021).

Several computational approaches have been developed to identify novel marker genes from scRNA-seq data in non-plant systems. The Seurat software first identifies highly variable genes across different clusters and then defines marker genes based on a statistical test between a cluster of cells against other clusters of cells (Butler et al., 2018). SCMarker is a statistical approach where marker genes are selected as mutually exclusively expressed with some other genes based on a mixture distribution model (Wang et al., 2019). A database called CellMarker was developed to provide a comprehensive overview of cell markers in both human and mouse (Zhang et al., 2019c). Using a traditional method called fisher linear discriminant analysis, scID selected marker genes for predefined cell clusters (Boufea et al., 2020), however, such method does not allow annotation of individual cells at intermediate developmental stages, thus is not very useful for plant research. In plants, ∼1500 cell-type marker genes have been determined from FACS-based gene expression data in the root cells of *Arabidopsis* (Birnbaum et al., 2005; Brady et al., 2007; Bruex et al., 2012; Efroni et al., 2015; Li et al., 2016). Recently, scRNA-seq data has been used to identify novel cell-type markers in plants by identifying genes with predominant expression in particular cell clusters with known identities (Jean-Baptiste et al., 2019; Shulse et al., 2019). However, complex heterogeneity of cell populations is sometimes characterized by multiple sub-populations of cells within one cluster, which limits the accurate identification of novel markers.

Machine learning (ML) has been widely applied to solve classification problems in genomics (Libbrecht and Noble, 2015). With regard to scRNA-seq data, supervised ML algorithms have been used to build cell-type classifiers (Alquicira-Hernandez et al., 2019; Pliner et al., 2019; Zhang et al., 2019a), which have outperformed traditional correlation-based approaches. However, none of these ML methods addresses the question of selecting marker genes in scRNAseq data (see **Table S1** for a comparison of 16 state-of-the-art machine learning methods for single cell type assignment.). Feature selection is a key component of modern machine learning methods because it provides interpretability to the ML models (Azodi et al., 2020). Feature selection refers to a class of techniques that assign scores to the input features to indicate how much each feature contributes to the performance of a predictive ML model (Cai et al., 2018). For example, a Support Vector Machine (SVM)-based recursive feature elimination was used to identify marker genes to differentiate developing neocortical cells from neural progenitor cells (Hu et al., 2016). These novel marker genes not only perform better than traditional gene sets, but also uncover hidden regulatory networks with novel interactions (Hu et al., 2016). We have also developed a feature selection-based approach to determine key regulators of transcription regulatory networks (Song et al., 2020).

In this study, we integrated five published scRNA-seq datasets from the *Arabidopsis* root containing over 25,000 cells and 17 cell clusters. We first compared seven machine learning methods for classification of ten different root cell types in *Arabidopsis*. We selected the best performing models, random forest (RF) and SVM, to use for the downstream analysis. For RF, we used a novel feature selection method called SHapley Additive exPlanations method (SHAP). Compared with traditional variable importance algorithms that only display results across all samples, the SHAP method allows us to calculate SHAP value for each observation, which greatly increases the model transparency (Lundberg et al., 2020). In comparison, we used the method suggested by Hu et al. (2016) to identify SVM-method based marker genes (SVMM). The SHAP and SVMM markers were compared to other sets of marker genes that have been published (KNOW), selected using correlation (CORR), from bulk RNA-seq (BULR), and those used in the index of cell identity model (ICIM).

We further demonstrated the power of machine learning based marker selection is not dependent on any specific cell type assignment approach. For example, we trained SPmarker on cells that were labeled by WEREWOLF (WER)-GRP promoter lines and identified new WER expressed cells. We also used SPmarker to determine annotations from additional markers that specify cell developmental stages in the root hair and epidermal cell types (Ryu et al., 2019). These new cell types could not be defined using traditional methods such as the ICI approach. As independent tests, the SPmarker method successfully assigned cells to respective cell types in two newly published data sets that were not used in the training of the models. We found that majority of new cell marker genes identified by SPmarker are not identified by traditional methods, and orthologous genes of SPmarkers showed significant overlapping with single cell marker genes found in rice, and in root hairs in five plant species, suggesting our approach can identify novel marker genes to facilitate cell type identification in scRNA-seq data from diverse plant species.

## Materials and Methods

### Data preprocessing

The scRNA-seq data of root cells from five publications were downloaded (Denyer et al., 2019; Jean-Baptiste et al., 2019; Ryu et al., 2019; Shulse et al., 2019; Zhang et al., 2019b). For each dataset, raw counts were used as input data, and any samples from mutant background or under treatment were removed. A gene was retained if it was expressed in more than three cells, and each cell was required to have at least 200 but no more than 5,000 expressed genes. The cells that have over 5% mitochondrial counts were removed. A global-scaling normalization method and multicanonical correlation analysis (Seurat v3.1) were used to normalize the expression data and to remove batch effects (Butler et al., 2018). Scrublet tool (Wolock et al., 2019) was used to predict doublet cells in this dataset. To correct for dataset-specific batch effects, multicanonical correlation analysis was used (Seurat v3.1). The normalized expression values in this merged dataset (57,333 cells and 29,929 genes) were used for the downstream training process. In processing of scRNA-seq data from a GFP-tagged line, raw reads were mapped to the TAIR10 reference genome using Cell Ranger pipeline (v2.1.1) with default settings (Zheng et al., 2017) to generate an expression matrix of 17,687 cells and 22,118 genes.

### Cell type annotation and training data preparation

To assign cell types to cells collected from the previous five datasets, Index of cell identity (ICI) score was computed (Efroni et al. 2015), for 15 root cell types include trichoblast, cortex, LRM (Lateral Root Meristem), Late_PPP (Late Phloem-Pole Pericycle), protophloem, meristematic xylem (Meri_Xylem), phloem_CC, protoxylem, phloem, pericycle, endodermis, atrichoblast, columella, QC, and Late_XPP (Xylem-Pole Pericycle). Cell type with the highest ICI score was assigned to the cell as final cell type label. To compare performance of our methods by using different ICI thresholds, we set two cut-offs: ICI > 0.5 or >0.91. There are 6,662 cells with ICI scores higher than 0.9 and only seven cell types were retained (> 100 cells).

Before the training process, two steps were conducted to balance the number of cells for each cell type: 1) Five cell types with small number of cells (<300) were removed; 2) 5000 cells from atrichoblast and cortex were randomly selected to reduce the size of the training sets. Finally, 25,618 cells from ten cell types were used for our analysis (Table S1). For GFP-tagged cells, we labeled 955 cells as ‘positive’ with reads mapped to WER-GFP gene, and other randomly selected 955 cells as ‘negative’ examples. Same analysis was performed using WER-AT (reads mapped to AT5G14750, WER gene) to label 1,970 cells as ‘positive’ and 1,970 cells as ‘negative’ examples. To train the models, the datasets were divided into a training dataset of 90% cells and an independent testing dataset of 10% cells. The training datasets were separated into sub-training (80%) and validation (20%) sets for five-fold cross-validation. The independent testing dataset was used to compare the performance of the machine learning methods.

### Machine learning classification

Machine learning approaches evaluated in this work include SVM, KNN, RF, baseline NN, triplet NN, contrastive NN (sklearn v 0.23.1 and Keras v2.2.4) (Pedregosa et al., 2011)(Chollet, 2015). All implementations of neural networks were modified based on a published study (Alavi et al., 2018). Although a few published methods were able to classify cell types, we did not find methods that can provide the flexibility to select marker genes using all these machine learning methods (see **Supplementary Table S1** for a discussion of a list of published methods). Therefore, instead of trying to compare to other stand-alone methods that implemented specialized methods for selecting marker genes, we use a generic machine learning package (sklearn) such that the methods are more directly comparable. Details for each machine learning approach are briefly described in **supplementary methods**. Source code for our SPmarker pipeline is available at github (https://github.com/LiLabAtVT/SPMarker).

## Results

### Comparison of different machine learning models for cell-type classification

The SPmarker pipeline includes two major steps (**Figure S1**). In the first step, the expression data of cells from different datasets were normalized and integrated using an established approach (Butler et al., 2018). The identities of these cells were assigned by some existing approaches such as: (1) using the ICI method (Efroni et al., 2015), (2) an internal GFP marker gene, or (3) by manually identified developmental stage related markers from a trajectory analysis (Ryu et al., 2019). Genes with highly variable expression were used as features for downstream analysis to select more specific marker genes (**Figure S1A**). In the second step, machine learning methods were trained and compared to determine methods that best predict cell types based on gene expression data. We selected the two best performing methods for follow up analysis to determine marker genes using feature selection (**Figure S1B**). The top 20 (default setting) genes with the highest feature importance in each cell type were selected as the marker genes.

Five scRNA-seq datasets of *Arabidopsis* roots were used in the training of the machine learning models (see Materials and Methods). The five data sets were merged into a single data set with 57,333 cells (**Figure S2**). To test different machine learning methods, we first used the ICI method to label the cell type for each cell (Efroni et al., 2015), followed by testing other methods of labeling cell types. The ICI score (0 <= score <= 1) of each cell represents a similarity of each cell to one of the 15 known cell types in *Arabidopsis* root and only 56% of all cells were able to be assigned by ICI (ICI > 0.5).

Seven machine learning methods were compared to determine which method performed best to classify cell types **(Figure 1)**. (Hoffer and Ailon, 2015)(Koch et al., 2015) (Gulli and Pal, 2017; Schmidhuber, 2015). These methods were selected because they represent approaches where each is based on distinct underlying mechanisms. The Area Under Precision-Recall Curve (AUPRC) values are shown for all the methods (**Figure 1A**), while other evaluation metrics were also calculated and compared between methods (**Figure S3 and S4**). The SVM and RF have highest AUPRC among the seven models (**Figure 1A**). For the deep learning-based models, the contrast NN, and triplet NN had similar performance but had higher AUPRC than the baseline NN. In general, non-neural network models showed relatively higher AUPRC than the deep learning-based models. This is not unexpected and further optimization of hyperparameters for NN models might improve the performance of NN based approaches. Performance comparisons were also obtained using seven other metrics (**Figure S3, S4, S5**). Tegardless of evaluation metrics used, the performances of SVM and RF are superior. Interestingly, machine learning models performed better for some cell types such as the trichoblast, atrichoblast, endodermis, and cortex, than other cell types such as the quiescent center (QC) cells (**Figure 1B**). This is not entirely due to the low number of cells in QC (537 cells, see **Table S2**) because other cell types such as phloem companion Cell (phloem_CC) and protophloem have similar or fewer cells (565 and 380 cells respectively) but the machine learning model performances on these cells are higher than QC (**Figure 1B**). Due to their better performances, RF and SVM were used to select marker genes using feature selection. Other methods were not further compared but can be tested if their performance can be improved using different hyperparameters.

**Figure 1.**
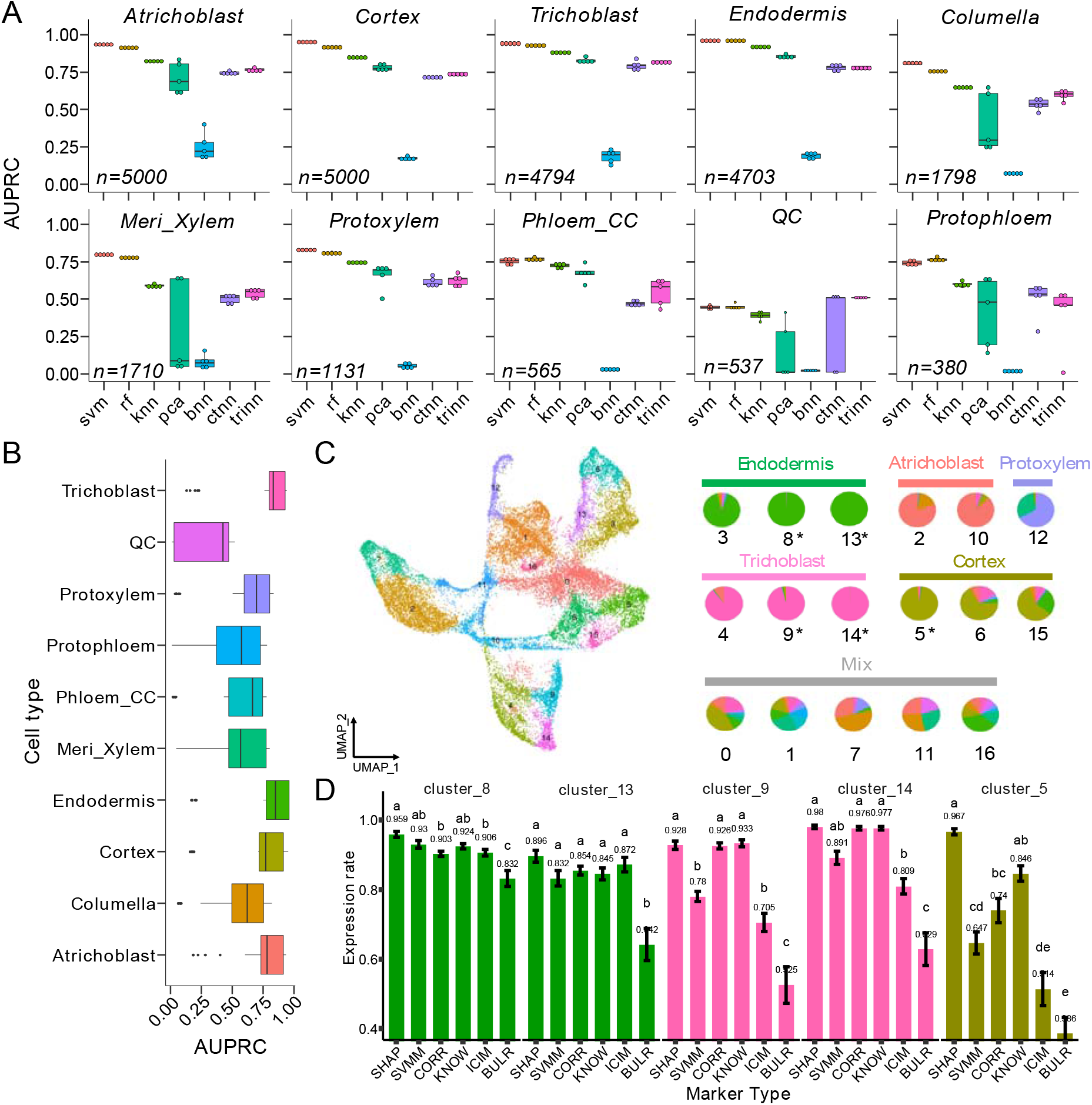
Classification performance of ten root cell types of *Arabidopsis*. **A**. Comparison of seven machine learning models on cell type classification. AUPRC means Area Under Precision-Recall Curve. svm: support vector machine. rf: random forest. knn: K-Nearest Neighbors. Pca: Principal Component Analysis. bnn, ctnn, and trinn: baseline, contrastive, and triplet neural networks. **B**. Comparison of classification performance of all the ten cell types. **C**. A UMAP plot where cells were clustered into 17 clusters. The right pie plot indicates cell composition in each cluster. If a cell type with cells occupies over 50% in a cluster, this type is defined as the dominant cell type. The labels above the pies are names of the dominant cell types of the clusters. Otherwise, the label for the clusters is ‘Mix’. * indicates clusters with more than 95% of cells belong to the same cell type. **D**. Comparisons of proportion of expressed cells among the six marker types. All pair wise comparisons are statistically significant as indicated by different letters (a, b, c, d, and e). If two bars have the same letter, then they are not significantly different from each other.

One way to compare the performance of marker genes is to compare their representations in different cell clusters in the scRNA-seq data. To make these methods comparable, we selected only the top 20 markers for each cell type for SHAP, SVMM and CORR respectively. We also selected 180 BULR, 161 KNOW and 232 ICIM markers. We cannot select exactly 200 markers for these published types of markers because they were predetermined by previous publications (see **Table S3** and **S4** for list of these marker genes). We have identified 17 clusters from the 25,618 single cells using Seurat and assigned cell type identities based on the ICI score for each cell. We focused on five clusters (5, 8, 9, 13, and 14) with a dominant cell type that accounts for over 90% cells in each cluster (**Figure 1C**, see **Figure S6** for a comparison of all cell clusters). These clusters were selected because they are the most homogenous clusters, making the results easier to interpret. For each marker gene in each cluster, we calculated the proportion of cells in which this marker gene is expressed, and we calculated the average “fraction of expressed cells” for all 20 markers in each cluster. For a more specific example, given 20 SHAP markers for cluster 8, on average each SHAP marker is detected in 95.9% of cells (**Figure 1D**, Cluster 8, Endodermis). In these homogenous cell clusters, the SHAP markers achieved a higher or similar proportion of the expressed cells as compared to all the other markers. In the cluster 5 that represents cortex, the SHAP markers had significantly (*p* < 0.05) higher expression rate (96.7%) than all other marker types (< 85%) (**Figure 1D**). BULR markers were detected in the lowest number of cells and followed by ICIM markers. Because the ICI method does not require all ICIM to be detected in a given cell, it is not surprising that ICIM were only found in 50% of cells in some clusters. On average, the percent of cells expressing SHAP markers is 29% more than ICIM and 67% more than BULR markers. Because both BULR and ICIM were determined using data not from scRNA-seq experiments, these results might suggest BULR and ICIM include cell type-specific, but relatively low-expressed genes that cannot be detected by scRNA-seq.

### Using newly developed markers to assign cell identity

Because only 20 marker genes were selected for each marker type for each cell type, we compared the expression patterns for these six types of markers using heatmap (**Figure S7** to **S12**). Markers identified by SHAP, CORR and ICIM have stronger cell type-specific expressions than KNOW, SVMM and BULR. To further quantify the specificity of the SHAP makers and other marker types, we calculated their cumulative correlation with specific cell types in the atrichoblast, trichoblast, and endodermis (**Figure 2A-C**). These three cell types were selected because all of these cell types have a higher number of cells than other cell types and these cell types are consistently identified by the majority of marker types. The cumulative correlation rates for SHAP markers are among the top three in all cell types, suggesting stronger preferential expression for this marker type. SHAP markers have similar performance as compared to CORR markers in atrichoblast and have higher performance than CORR markers in two other cell types. ICIM markers are among the top three in atrichoblast and endodermis but ranked fourth in trichoblast. This is consistent with the observation that not all ICIM markers were detected in all cells.

**Figure 2.**
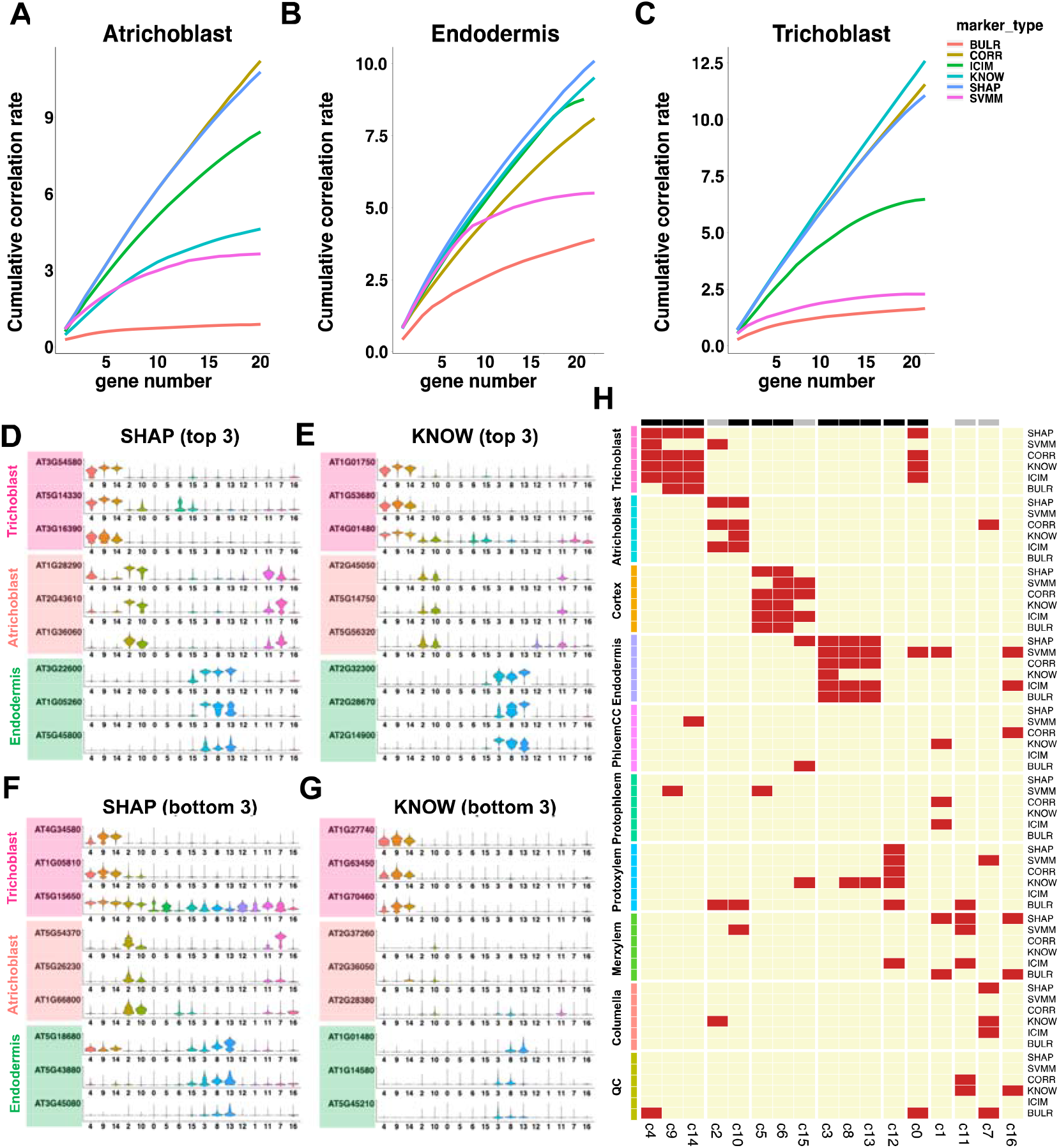
Assignments of cell identity using newly developed markers. **A-C**. Cumulative correlation plot for top 20 markers for six types of markers. **D-G**. Violin plots that show the expression of top three markers (**D, E**) and bottom three markers (**F**,**G**) in three cell types, across all clusters. **H**. Heat map of cell types assigned to each cluster by different markers. On top of the heatmap, black bars show clusters assigned consistently by four or more methods, and grey bars show clusters assigned consistently by three out of six methods.

To demonstrate the specificity for SHAP markers, we plotted the top three most specific markers from the 20 selected SHAP and KNOW markers (**Figure 2D** and **E**). We also plotted the expression of the bottom 3 markers (ranked 18, 19, and 20 by marker specificity) from the 20 markers (**Figure 2F** and **G**). We found that the SHAP markers showed high cell-type specificity in both cases whereas the specificity of KNOW markers is lower in at least 7 cases for those ranked at 18 to 20. One interesting observation is that, although SVMM markers do not show high cumulative correlation, most of these markers are highly expressed in multiple clusters (**Figure S13**), suggesting SVM provides a different approach to detect cell types.

One potential limitation of machine learning models selected markers is that other model parameters such as the decision thresholds for RF and feature weights for SVM associated with each group of markers has to be evaluated using model-dependent algorithms. Because of the high correlation of SHAP markers with specific cell types, we develop a voting procedure to simplify the process of assigning cell identities using newly developed markers as well as existing marker genes (see method section, **Figure 2H**). Using this method to the 17 clusters, we found 15 clusters were assigned consistently to the same cell types by three or more marker types (black and grey color marked the clusters in **Figure 2H**). These results showed that cluster assignments are consistent between existing markers and new marker genes.

### Identification of marker genes with different training labels

The SPmarker method is more flexible than traditional approaches because our method can be trained on different cell labels and select different sets of marker genes to classify cell types. To compare our method with traditional methods, we tested three different scenarios: (1) label cells with a different ICI threshold; (2) label cells in the same lineage under different developmental stages; and (3) label cells with an internal GFP marker.

We first compared the performance of SPmarker by using two different ICI thresholds, 0.5 and 0.9 (**Figure 3A and 3B**). Marker genes were selected by RF and SVM models and only the top 20 marker genes were used for the analysis to match the number of genes in ICIM and other marker types (CORR, KNOW, and BULR). Random forest models were trained using these marker genes separately and performances were compared using AUPRC. We have only five cell types with enough cells or enough marker genes from all methods for comparison with ICI > 0.9 (see **Figure S14** for all five cell types). ICIM performed best in both thresholds in five cell types tested, which is expected because the ICIM is used to label cells. Interestingly, we found that the performance of SHAP and SVMM markers increased significantly for prediction ICI>0.9 cells as compared to ICI>0.5 cells. These results may suggest that ICI0.9 cells are more specific as compared to cells with ICI > 0.5 and they are easier to be classified using different sets of markers. In contrast, CORR, KNOW and BULR markers did not show improvement of performance, partly because these marker genes were determined not based on cells labeled by training samples thus are less flexible than SPmarkers.

**Figure 3.**
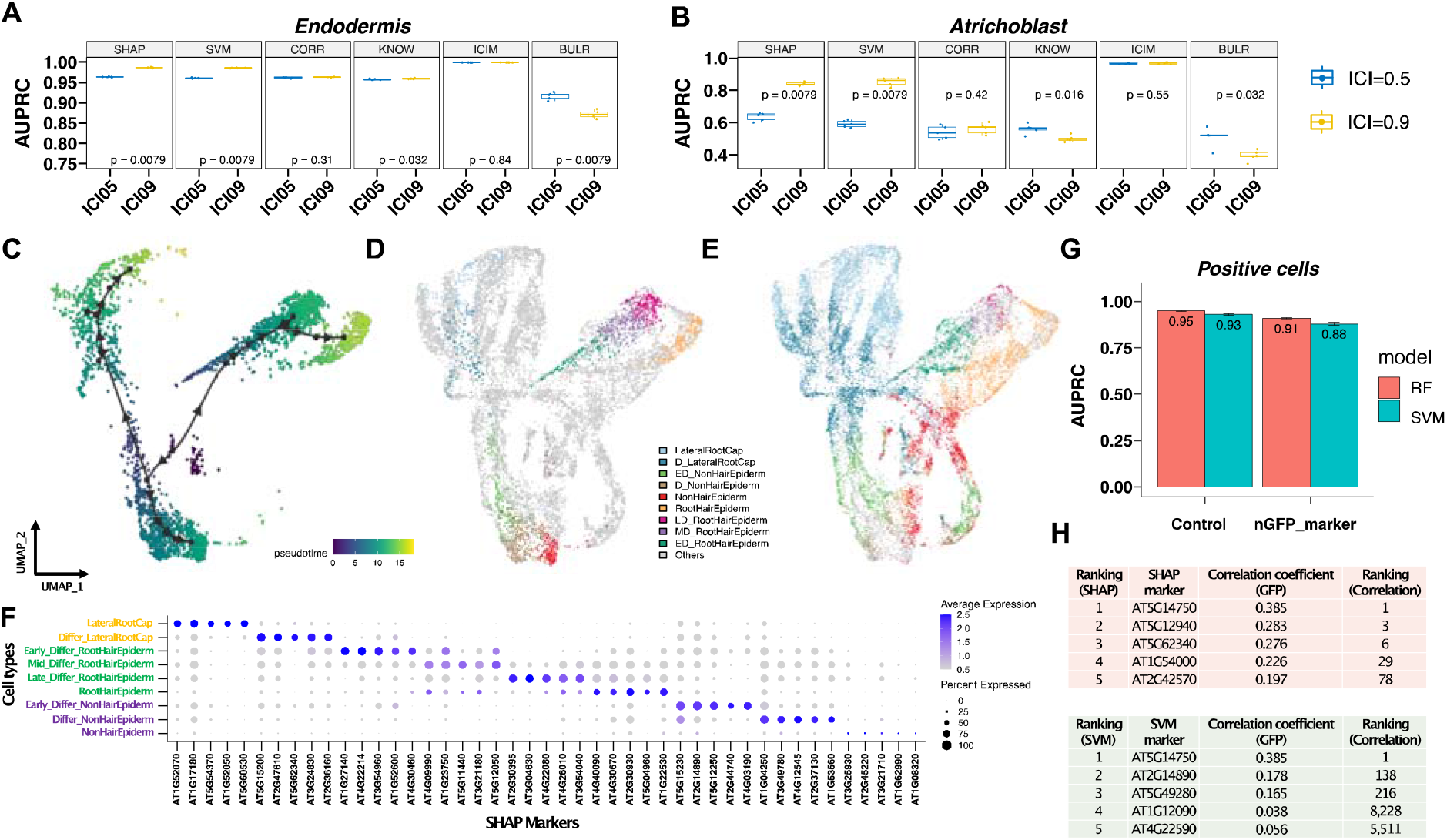
Identification of marker genes with three different ways to label cells. **A-B** Comparison of classification performance based on ICI labeling method between 0.5 and 0.9 thresholds in endodermis **(A)** and atrichoblast **(B)** cell types. p value < 0.05 indicates significant differences between ICI05 and ICI09 groups. **C-F**. Label cells under different developmental stages. **C**. Trajectory analysis of cells from root hair, non-root hair and lateral root caps from our published single cell data. **D**. A UMAP of nine developmental stages derived from these three cell types. ‘Others’ label indicates cells from other publications including four studies with trichoblast and atrichoblast cells and with WER cells generated from a WER-promoter-GFP line (Ryu et al., 2019). In the legend, letter ‘D’ means ‘Fully Differentiated’, ‘ED’ means ‘Early Differentiated’, ‘MD’ means ‘Middle Differentiated’, ‘LD’ means ‘Late Differentiated’. **E**. A UMAP shows predictions of cell identities from the other publications. ‘Others’ label indicates cells from Ryu’s study (2019). **F**. Identification of SHAP markers in the nine developmental stages. **G-H**. Identification of markers with labeling cells using internal GFP marker. **G**. Comparison of classification performance on GFP-labeled WER cells (positive cells) between using all genes (control) and genes without GFP marker (nGFP_marker) for both RF and SVM models. **H**. Ranking of best SHAP and SVM markers to predict WER-GFP positive cells. The third column indicates expression correlation between GFP and other genes. The fourth column indicates ranking based on correlation values.

We next tested whether we can use machine learning to transfer labels from one experiment to another (**Figure 3C-F**). First, we used our published single cell data (Ryu et al., 2019) and selected cells from root hair, non-root hair and lateral root caps. These cell types were selected because they are all epidermal cells and represent distinct cellular functions. These cells were further classified into nine different developmental stages based on the analysis of trajectories (Ryu et al., 2019). SPmarker was trained on this data to select SHAP and SVMM markers and predictions were made on the other four datasets in the integrated root cell map (**Figure 1C**). In the UMAP plots, we found that cell types from Ryu (2019) follow three separate trajectories that correspond to the three selected cell types (**Figure 3C**). Most importantly, the labeled cells (**Figure 3D**) overlap strongly with the cells from other publications and form similar trajectories. For example, from Ryu (2019) data (**Figure 3C** and **3D**), we found that the ‘differentiating lateral root cap’ are located close to the center of the UMAP (dark blue), whereas the mature lateral root cap cells are located towards the outskirt of the UMAP plot (light blue). Cells from other four publications show similar distribution (**Figure 3E**, dark and light blue cells). This is also observed for the root hair lineages (dark green, purple, and dark pink cells), and for the non-hair lineages (light green, brown, red cells). We also identified new marker genes from these cell types at different developmental stages which showed stage specific expression patterns (**Figure 3F**). These results show that SPmarker can identify cell types and marker genes with fine-grained resolution for cell types at separate differentiation stages.

Interestingly, there are some cells that were not classified to the same type using RF or SVM (**Figure S15**). In these cells, RF predicts these cells as non-hair epidermal whereas SVM predict these cells as lateral root caps. These cell types are relatively similar and therefore, the prediction actually reflects the biological similarity of these cells and intrinsic ambiguity of cell identities of these cells.

Finally, we tested our method using internal control genes as labels **(Figure 3G and H)**. In our published scRNA-seq data (Ryu et al., 2019), we used a WER-promoter-GFP line to generate scRNA-seq data. WER-promoter-GFP was highly expressed in root cap, moderately expressed in artrichoblasts. Therefore, in the single cell gene expression matrix, we were able to identify reads that mapped to both GFP (WER-GFP) and the WER (WER-AT, AT5G14750) genes. We labeled cells using reads mapped to WER-GFP first and trained the model to predict WER-GFP positive cells. WER-GRP was removed from the training data such that this gene will not be used as marker itself. The same analysis was performed using WER-AT. As expected, when we labeled cells by WER-GFP and select marker genes using either RF or SVM, the best marker to predict WER-GFP positive cells are WER-AT genes in both methods (**Figure 3H**). More interestingly, when we removed both WER-GFP and WER-AT from the gene expression matrix, we can still predict WER positive and negative cells with high AuPRC (**Figure 3G**, nGFP_marker), suggesting that other marker genes can also provide prediction to the WER-positive cells. We further compared the correlation of SHAP and SVMM markers to the expression of the internal WER-GFP tag. We found that the correlation is low (r = 0.385) and most importantly, the top ranked genes by SHAP or SVMM are not top ranked genes by correlation. When we labeled with WER-AT instead of WER-GFP we observed similar results but obtained a different list of high-ranking genes (**Figure S16**). Since WER-labeled cells were not included in the original ICI cell types, these results demonstrate that our machine learning method is applicable in identifying new cell types with alternative methods of cell labels.

### Most machine learning derived markers are new markers

Because SHAP and SVMM are very different from correlation markers in the WER-GFP analysis, we sought to understand how many new markers can be identified in other cell types. We identified SHAP marker genes that are unique to each cell type using ICI > 0.5. We found 1,840 and 1,460 unique marker genes for cortex and atrichoblast respectively, while 63 and 37 unique marker genes were found for the QC and protophloem (**Figure 4A**). In other words, there are almost 30 times more unique SHAP marker genes in cortex as compared to QC cells. By overlapping the cumulative *SHAP* values from genes with unique SHAP marker, we found fewer than 50 unique genes accounted for 50% of the total *SHAP* values in each cell type (**Figure S17**). This suggests that QC has a small number (**Figure 4A**, 63 genes) of unique SHAP markers and more markers (**Figure S17**, 475 genes) are shared with other cell types, whereas cortex or atrichoblast have large numbers of unique SHAP markers but a small fraction of these markers carry the most weight.

**Figure 4.**
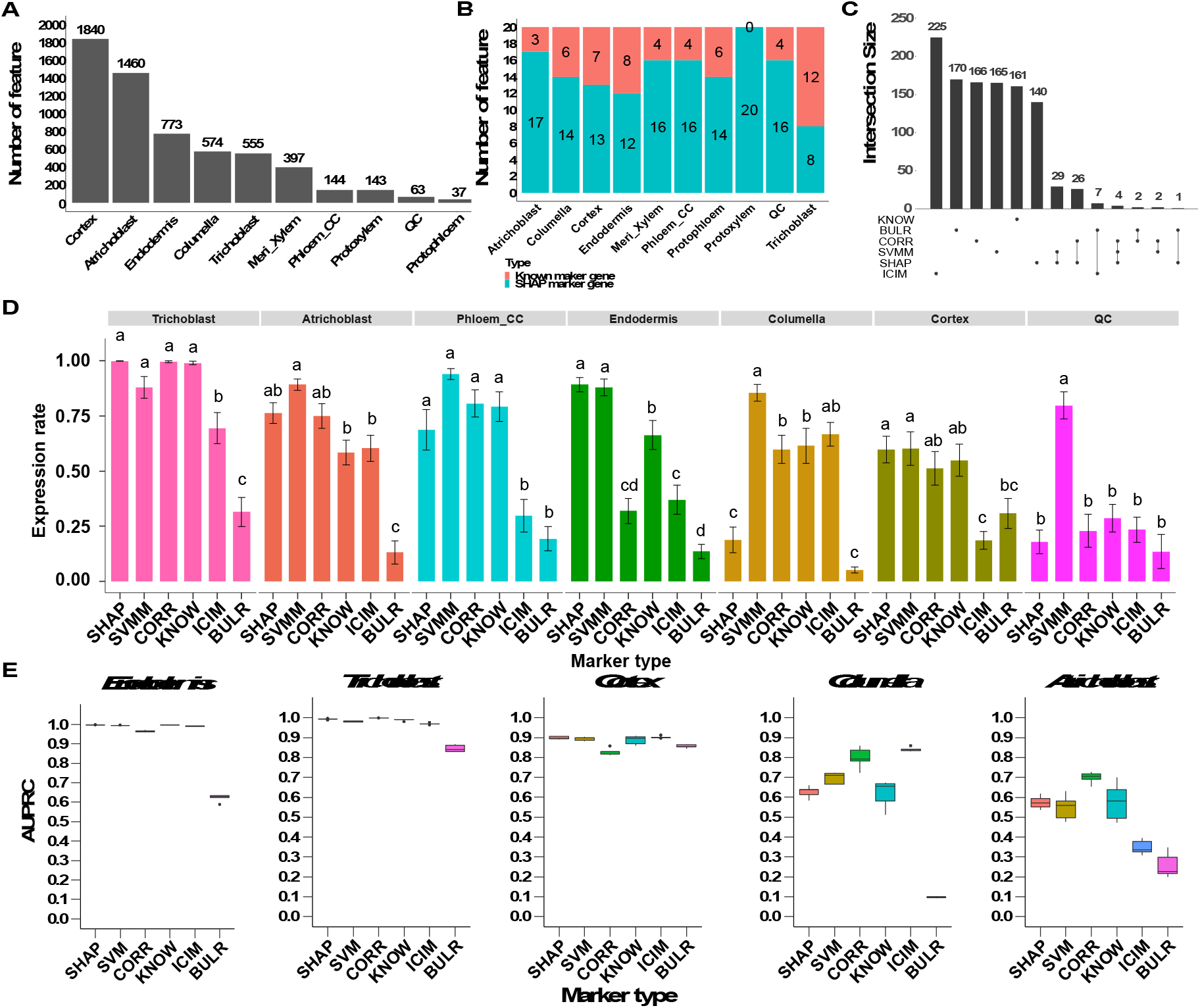
Testing of new markers with independently labeled cell types. **A**. Number of SHAP marker identified in each cell type. **B**. Comparison of number of SHAP and known markers in the 20 genes with the highest SHAP value in each cell type. **C**. Summary of gene counts from six marker types. Set size means gene count of different marker types. The dots under the bars mean the genes specifically exist in the relative marker type. The line connected between two or more dots under the bars mean genes exist in two or more marker types. If two or more marker types do not have connection, it means these groups do not have shared genes. **D**. Comparisons of proportion of expressed cells among the six marker types. All pair wise comparisons are statistically significant as indicated by different letters (a, b, and c). If two bars have the same letter, then they are not significantly different from each other. **E**. Comparison of six marker types on cell type classification.

To study the identity of the SHAP marker genes, we focused on the top 20 genes in each cell type with the highest SHAP value. There are 146 genes out of 200 SHAP marker genes (73%) that were not identified before in a collection of 1,813 marker genes from other publications (**Table S6**). In particular, in the Protoxylem, all SHAP markers are new (**Figure 4B**) and 80% of SHAP markers from the atrichoblast and phloem_CC are new markers (**Figure S18)**. The same results were observed for SVMM, where the majority of new markers were specifically found by SVM method but not by other methods (**Figure S19**). Interestingly, there is little overlap between these marker genes such that 93.5% or 1,027 marker genes are unique to a single method. When we compared the top 200 marker genes identified by six different methods, we found that most markers were found by a single method, whereas only 71 markers were found by two methods (**Figure 4C**) and there is no single marker ranked as top 20 by more than three methods. We also found unique biological functions of these newly identified marker genes (**Figure S20, Table S7, S8 and S9**).

To validate these newly identified marker genes, we searched literature for published orthogonal experimental evidence for cell type specificity of these new marker genes. We have identified 11 cases of published wet-bench data supporting our new markers, and all the cases were not found by traditional methods for single cell data analysis (**Table S10, Table S4, and supplementary methods**). More importantly, these 11 cases were generated by independent publications and thus are unbiased validation of our newly identified markers.

### Testing new marker genes with independently labeled cell types and new single cell data

When we first implemented SPMarker, only five single cell datasets were available from *Arabidopsis* roots. To test whether SPmarker can predict cell types in data that were not used in training, we evaluated the model performance using two newly generated single cell datasets for *Arabidopsis* roots (**Figure 4D and 4E, and Figure S21**). In both datasets, different approaches were used to label cell clusters. In one paper, the top variable genes from each single cell cluster were selected and correlation with published bulk-RNAseq data were used to assign cell types (Wendrich et al., 2020). In the other paper, three different methods were used to assign cell types and the consensus of three methods were used for final label of cell types (Shahan et al., 2020). To evaluate the new marker genes identified in our approach and compare them to other existing marker genes, we calculated how many cells in each cell type have these marker genes expressed (**Figure 4D, and Figure S20**). Only 7 or 6 cell types were exactly matched between our cell names and these two newly published papers. For example, for cells identified as trichoblast in one paper (Wendrich et al., 2020), more than 99% of cells also expressed marker genes from SHAP, CORR and KNOW categories. 88.1% of cells expressed SVMM markers, and fewer than 70% of cells expressed ICIM or BULR markers. This result shows that if only ICIM were used to assign cell types, ∼30% of cells will not be assigned due to the lack of marker genes. This also shows that SHAP, CORR and KNOW markers, and to a lesser extent, SVMM, work well to define trichoblast cells determined in this publication.

Not all machine learning based markers performed well, for example, SHAP markers are not performing well in columella and QC cells. Interestingly, the BULR markers showed lower performance in all cell types, one possible reason is that BULR markers are lowly expressed and cannot be detected in single cell data, but they are highly specific to individual cell types. These results are also observed when evaluating these marker genes in the Shahan’s paper (2020) (**Figure S21**). Finally, we test the performance of different marker genes when we use them with a RF model to label cell types (**Figure 4E**). These models were trained using integrated dataset (**Figure 1C**) that does not include the new data thus serve as a completely independent validation. We found that AUPRC is close to 1.0 for five marker types in endodermis and trichoblast, and close to 0.9 for cortex, representing high performance of the model to transfer annotation to a different dataset. In contrast, AUPRC for all types of markers are more variable and lower for columella and atrichoblast, suggesting additional methods would be needed to determine the best way to assign cell types of cross studies in these cases. CORR has better performances in two cell types where the overall AUPRC are lower than 0.9, suggesting when cells are harder to classify, CORR might have better performance.

### Newly identified marker genes can be used to assign cell types in other species

As scRNA-seq experiments expand to include non-model plant species that lack cell-type markers, it becomes challenging to accurately determine cell types in these species. A major usage of new marker genes is to expand the list of candidate marker genes in other species, because some marker genes in *Arabidopsis* may have altered their functions in other species or may be absent from genomes of other species. To this end, we compared the marker genes identified from a newly published, single cell sequencing data from rice roots (**Figure 5A and Table 1**) (Liu et al., 2021). There are only 3 clusters that were given the same cell type names in both the *Arabidopsis* and rice data; therefore, we can only compare marker genes in theses 3 clusters. We found very small number of *Arabidopsis* marker genes whose orthologous genes were also found in corresponding cell types in rice. For example, for cortex, there are no marker genes from the ICIM and BULR that were also found in rice cortex cluster (**Table 1**). There are only 10 KNOW markers were also found in rice cortex cluster, and such overlapping is not statistically significant. Although ICIM and KNOW marker showed a significant overlapping in endodermis and trichoblast respectively, the absolute number of overlapping markers are small, which was also found by the original publication (Liu et al., 2021). When we compared the SPmarkers (SHAP and SVMM), we found substantial increase of overlapping markers and such overlapping are statistically significant in five out of six cases (**Table 1**). In contrast, the overlapping of CORR markers is only significant for trichoblast (p < 0.01) but not for other two cell types. When we analyzed how many markers were also detected in the three cell types (**Figure 5A**), we found approximately 60% of cortex and endodermis cells and more than 75% of trichoblast cells from rice also have SPmarkers. CORR markers showed lower detection rates than one or both of the SPmarkers in two cell types. ICIM and KNOW markers were missing from one cell type and showed similar or lower detection rates than SPmarkers.

**Figure 5.**
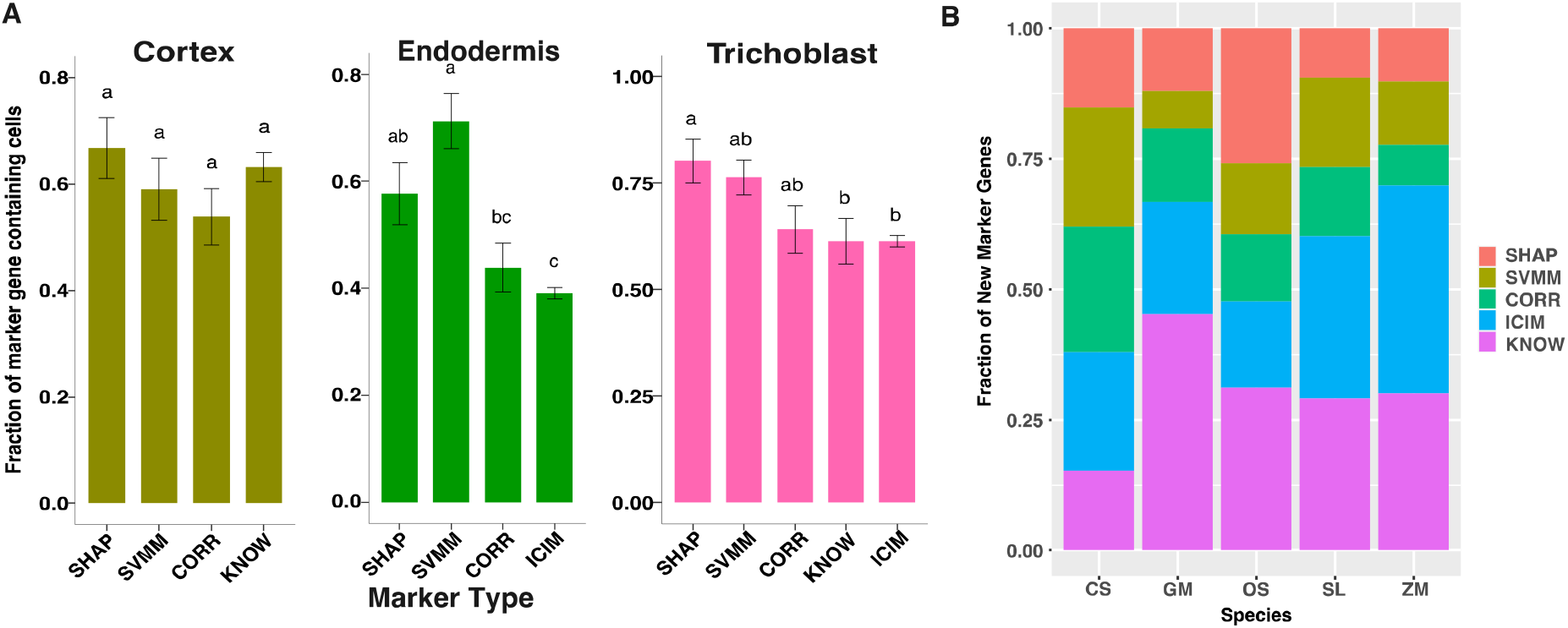
Testing of new identified markers in other species. **A**. Comparisons of proportion of expressed cells among the four marker types in a published rice scRNA-seq data (Liu et al., 2021). All pair wise comparisons are statistically significant as indicated by different letters (a, b, and c). If two bars have the same letter, then they are not significantly different from each other. **B**. Number of root hair marker genes identified using five marker types in five species including *Cucumis sativus* (Cs), *Glycine max* (Gm), *Oryza sativa* (Os), *Solanum lycopersicum* (Sl), and *Zea mays* (Zm).

**Table 1.**
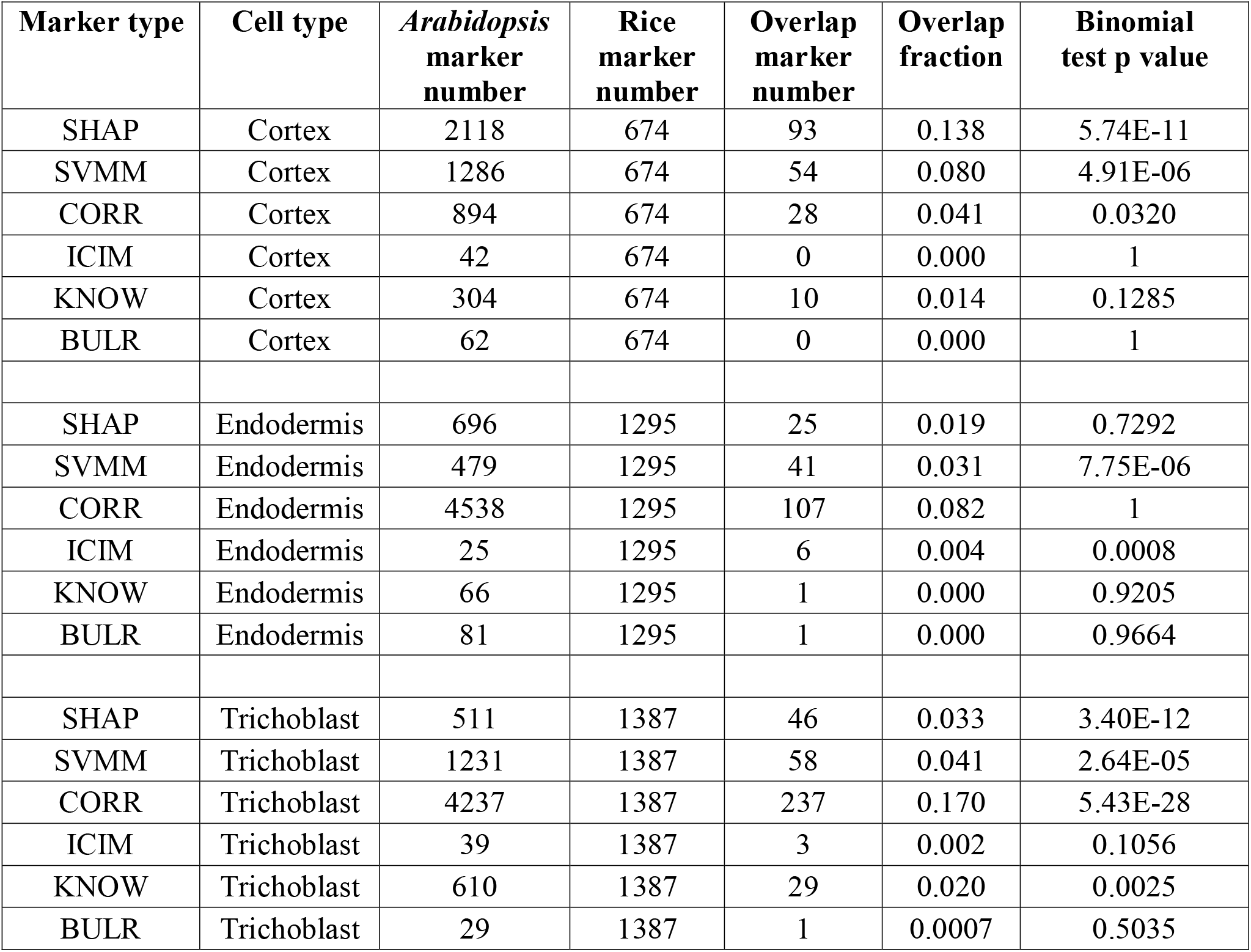
Comparison of number of overlapped rice markers in three cell types among six marker types. *Arabidopsis* marker number is the number of orthologous genes of *Arabidopsis* markers in the rice genome according to Phytozome annotation. Overlap ratio is calculated as overlap marker number divided by rice marker number. Binomial test p values were calculated using one hundred random draws of same number of marker genes and compared these random overlap ratios to the observed overlap ratio.

| Marker type | Cell type | *Arabidopsis*<br>marker<br>number | Rice<br>marker<br>number | Overlap<br>marker<br>number | Overlap<br>fraction | Binomial<br>test p value |
| --- | --- | --- | --- | --- | --- | --- |
| SHAP | Cortex | 2118 | 674 | 93 | 0.138 | 5.74E-11 |
| SVMM | Cortex | 1286 | 674 | 54 | 0.080 | 4.91E-06 |
| CORR | Cortex | 894 | 674 | 28 | 0.041 | 0.0320 |
| ICIM | Cortex | 42 | 674 | 0 | 0.000 | 1 |
| KNOW | Cortex | 304 | 674 | 10 | 0.014 | 0.1285 |
| BULR | Cortex | 62 | 674 | 0 | 0.000 | 1 |
| SHAP | Endodermis | 696 | 1295 | 25 | 0.019 | 0.7292 |
| SVMM | Endodermis | 479 | 1295 | 41 | 0.031 | 7.75E-06 |
| CORR | Endodermis | 4538 | 1295 | 107 | 0.082 | 1 |
| ICIM | Endodermis | 25 | 1295 | 6 | 0.004 | 0.0008 |
| KNOW | Endodermis | 66 | 1295 | 1 | 0.000 | 0.9205 |
| BULR | Endodermis | 81 | 1295 | 1 | 0.000 | 0.9664 |
| SHAP | Trichoblast | 511 | 1387 | 46 | 0.033 | 3.40E-12 |
| SVMM | Trichoblast | 1231 | 1387 | 58 | 0.041 | 2.64E-05 |
| CORR | Trichoblast | 4237 | 1387 | 237 | 0.170 | 5.43E-28 |
| ICIM | Trichoblast | 39 | 1387 | 3 | 0.002 | 0.1056 |
| KNOW | Trichoblast | 610 | 1387 | 29 | 0.020 | 0.0025 |
| BULR | Trichoblast | 29 | 1387 | 1 | 0.0007 | 0.5035 |

Because we found a substantial increase of marker genes when compared to rice single cell data, we asked whether such comparison can be expanded to other species. However, no single cell data from other species are available at the time of this work. Therefore, we analyzed root hair cell expression from five plant species (cucumber, soybean, rice, tomato, and maize) where root hair specific expression data are available (Huang et al., 2017). To consider the most specific markers, we tested the top 20 markers from each marker type from single cell data. In these five species, we found a total of 172 genes are significantly differentially expressed in root hair cells that are also orthologous genes to the six types of marker genes (**Figure 5B**). SHAP and SVMM accounted for 26.1 to 60.7% of these root hair genes and these new marker genes increase the number of candidate marker genes by 35.3% to 154.5% in these five species.

## Discussion

The scRNA-seq technology provides a novel platform to analyze the transcriptomic profile of individual cells to characterize heterogenous cell populations in detail. In plants, this process is heavily reliant on the use of marker genes that are preferentially expressed in specific cell types. Here we introduce a machine learning based approach, SPmarker, to identify marker genes by analyzing their feature importance. Machine learning methods provide a number of principled approaches to evaluate marker performance including cross-validation, leave-out testing sets, and evaluation metrics such as auROC, auPRC and F1 scores. These evaluations methods allow us to compare different marker genes in a more rigorous and unbiased fashion. In addition, Seurat package is a standard method that needs prior knowledge of cell clustering to identify marker genes. We have compared marker genes identified by machine learning methods with those identified by FindAllMarkers function (with default parameters). We have found there is no statistically significant difference in the expression rate for SHAP markers as compared to Seurat identified markers (**Figure S23**). More importantly, the majority of SHAP and SVMM markers are different from Seurat markers (**Figure S24**). We have shown that these new markers can yield high performance using machine learning-based evaluation metrics. More importantly, because these machine-learning derived markers are not based on prior knowledge of gene functions, these markers may have new biological functions that are not characterized before.

In the evaluation of different machine learning methods, the SVM and RF methods outperform the three deep learning models (**Figure 2**). One possible reason is SVM and RF are effective for relatively small datasets or fewer outliers (Ali et al., 2012; Ben-Hur and Weston, 2010). The deep learning algorithms usually require relatively large dataset to work well and achieve good performance for solving more complex problems such as image classification (Zou et al., 2019). A previous study utilized the contrastive NN and triplet NN to successfully classify cells in mouse by using more than 100,000 cells to train these two models (Alavi et al., 2018), while our study used less than 30,000 cells. If more cells with accurate cell identity were available in *Arabidopsis*, the performance of the deep learning model in our study may be improved (Eraslan et al., 2019).

Cell populations of *Arabidopsis* roots are characterized by a high level of heterogeneity. Results from animal systems have demonstrated that, even within a cell population, the cells are not homogeneous because sub-populations may exist (Liu and Trapnell, 2016). Furthermore, it is not clear whether all cell types have been discovered for the *Arabidopsis* root (Zhang et al., 2019b), in particular, for cells in a transition stage or regulated by periodical signals (Voß et al., 2015). This highlights the importance of identifying new marker genes which may be expressed at different levels in sub-populations as compared to traditional marker genes derived from bulk RNA-seq and microarray experiments. However, traditional approaches for marker gene identification usually involve manual inspection of the cell population structure, which could be arbitrary (Luo et al., 2015; Usoskin et al., 2015).

Identification of new marker genes is particularly important for the research community of plant biology because cell type markers are largely unknown from non-model species. We have demonstrated that our machine learning based approaches can substantially expand the number of known root hair marker genes and that orthologs of these marker genes can also be found in other plant species. One future direction is to define root cell types in non-model species from cross-species mapping of marker genes and their expression pattern in roots.

## Author Contributions and Acknowledgments

HY, JL, QS and SL designed the experiments and performed the computational analysis. BZ and QL provided experimental validation of candidate genes. HY, JS and SL wrote the manuscript. We would like to acknowledge Jeffress Trust Awards Program in Interdisciplinary Research (to J.L., and S.L.); United States Department of Energy Funding (DE-SC0020358) to S.L., and J. S.; Hatch Programs from United States Department of Agriculture (to S.L.).

## Notes

### Competing Interest Statement

The authors have declared no competing interest.

### Summary of Updates

We have made substantial update to the methodology used in this article. Now the method can be trained on different types of cell labels, including (1) assign cell types based on cells that were labeled using published methods, (2) project cell types identified by trajectory analysis from one dataset to other datasets, and (3) assign cell types based on internal GFP markers.

